# CMV Replication Drives IFNγ-Mediated Sensitization of AML Cells to Cytotoxic Killing Through the NKG2C-HLA-E Axis

**DOI:** 10.64898/2026.04.24.720601

**Authors:** Wiebke Moskorz, Ron Patrick Cadeddu, Markus Uhrberg, Paul Sebastian Jäger, Ralf Grutza, Ramona Grothmann, Mirko Trilling, Sascha Dietrich, Christine Cosmovici, Rainer Haas, Jörg Timm

**Author notes:** Corresponding author: Wiebke Moskorz, Institute of Virology, University Hospital Düsseldorf, Heinrich-Heine-University Düsseldorf, Building No.: 22.21.02.26, Universitätsstr. 1, 40225 Düsseldorf. Ramona Grothmann, Clinic for Haematology and Stem Cell Transplantation, University Hospital Essen, Essen, Germany.

## Abstract

Human cytomegalovirus (CMV) infection represents a significant risk factor for transplant recipients, including patients undergoing hematopoietic stem cell transplantation (HSCT). Interestingly, several studies have reported an association between early CMV reactivation and a reduced risk of leukemia relapse, particularly in acute myeloid leukemia (AML). Given that CMV profoundly shapes the natural killer (NK) cell compartment, a contribution of CMV-primed NK cells to this effect has been proposed.

To explore this mechanism, we analyzed the relationship between NK cell functionality and CMV reactivation in the context of AML. Consistent with observations in peripheral blood, CMV-seropositive HSCT recipients displayed expanded NKG2C^pos^ NK cell populations within the bone marrow, characterized by high Granzyme B expression. CMV replication was associated with elevated plasma IFNγ levels, which *in vitro* rendered AML cells more susceptible to apoptosis when co-cultured with peripheral blood mononuclear cells. Importantly, IFNγ treatment modulated NK cell responses by inducing a variety of NK cell ligands including HLA-E on primary bone marrow-derived blasts and AML cell lines. In line with this, the activation of CMV-associated NKG2C^pos^ NK cells was enhanced upon stimulation with IFNγ-pretreated AML cells.

In summary, our findings demonstrate that CMV replication induces a transient increase in IFNγ levels that influences both AML and NK cells, ultimately enhancing AML cell susceptibility to NK cell-mediated cytotoxicity initiated through the NKG2C-HLA-E axis.

**Importance:** Previous studies suggested that CMV reactivation after HSCT may reduce leukemia relapse in AML patients, but the underlying mechanism remained unclear. Here, we show that CMV replication induces IFNγ release, which sensitizes AML cells to NK cell-mediated killing. This effect involves upregulation of HLA-E on AML cells and activation of expanded NKG2C^pos^ NK cells within the bone marrow. Our findings uncover a novel IFNγ-dependent link between CMV replication and enhanced NK cell cytotoxicity in AML, suggesting that combining IFNγ treatment with NK cell-based immunotherapy or NKG2A blockade could reduce post-HSCT relapse, even in CMV-negative patients.

## Introduction

Cytomegalovirus (CMV) infection is highly prevalent especially in older adults, a population with an increased risk for hematologic malignancies. Among patients undergoing hematopoietic stem cell transplantation (HSCT), CMV represents a significant clinical challenge; as CMV seropositivity alone has been associated with adverse post-transplant outcomes [1, 2]. Reactivation of CMV in immunocompromised HSCT recipients can cause severe complications, including CMV pneumonia, encephalitis and gastrointestinal disease. Intriguingly, several studies have reported a counterintuitive association between early CMV replication after HSCT and a reduced risk of leukemic relapse in acute myeloid leukemia (AML), myelodysplastic syndromes (MDS), chronic myeloid leukemia (CML) and acute lymphoblastic leukemia (ALL). The underlying mechanisms are still a matter of debate, while some contributing transplantation-related factors such as the type of conditioning regimen and the use of antithymocyte globulin (ATG) administration could be identified [3-11].

CMV infection induces long-lasting alterations in the immune system [12], which may enhance the clearance of residual leukemic stem cells thereby contributing to a lower relapse risk. In AML, NK cells display a skewed receptor repertoire and reduced cytotoxicity (reviewed in [13]), suggesting impaired NK cell function. Since NK cells are the first lymphocytes to reconstitute after HSCT [14, 15], they are among the earliest effectors capable of targeting residual leukemia as well as pathogens. CMV infection has been shown to enhance NK cell activity in vitro [16] and to drive a pronounced expansion of CMV-specific CD8 T cells, usually 5-10 % but occasionally up to 30-50 % of total CD8 T cells [17, 18]. In parallel, CMV infection promotes the expansion of NKG2C^pos^ NK cells (reviewed in [19]), which exhibit potent cytotoxicity, elevated IFNγ production [20, 21] and strong antibody-dependent cellular cytotoxicity (ADCC) [22, 23]. These NKG2C^pos^ NK cells have been proposed to mediate anti-leukemic effect observed in CMV-reactivating patients (reviewed in [24]). Despite the association between CMV reactivation and lower relapse rates in AML, most studies that do not stratify based on CMV peak titers report no improvement, or even a decline in overall survival, primarily due to increased non-relapse mortality [4, 5, 7, 8, 25, 26].

In this study, we investigated how CMV replication influences NK cell functionality in HSCT recipients, aiming to leverage its protective anti-leukemia potential while minimizing CMV-related risks. Our findings reveal that CMV replication induces a transient increase in IFNγ levels, which drives upregulation of HLA-E on AML cells. This renders the AML cells more susceptible to recognition and killing by NKG2C^pos^ NK cells through enhanced mobilization of Granzyme B and Perforin. These results suggest a mechanistic link between CMV-induced IFNγ signaling and NK cell-mediated leukemia control, offering a potential avenue for therapeutic exploitation post-HSCT.

## Material & Methods

### Study subjects

White blood cells from bone marrow (BM) and/or peripheral blood (PB) of patients with primary hematologic malignancies, including HSCT recipients and non-transplanted patients, were obtained through the Department of Hematology, Oncology and Clinical Immunology at the same institution under Ethics Committee approval (study number 2020-1222). Most of HSCT recipients received Mycophenolat-Mofetil combined with Tracrolimus as Graft versus host disease prophylaxis. Further patient characteristics are detailed in Supplementary Table 3. Peripheral blood mononuclear cells (PBMCs) from healthy blood donors were isolated from buffy coats provided by the Center for Blood Donation at University Hospital Düsseldorf. CMV serostatus was assessed using the LIAISON® XL system (DiaSorin) and plasma CMV DNA levels were quantified with the Cobas® 6800 system (Roche).

### Sample preparation

PBMCs from healthy blood donors were isolated by density gradient centrifugation and were stored at -196 °C until further usage. White blood cells from peripheral blood and/or bone marrow aspirates from patients with a primary hematologic disease were isolated by red blood cell lysis using isotonic lysis puffer (155 mM NH_4_Cl, 10 mM KHCO_3_ and 0.1 mM EDTA, pH 7.4). In short, peripheral blood or bone marrow was incubated with red blood cell lysis buffer for 10 min at a ratio of 1 to 10, centrifuged for 5 min at 500 g and washed twice with DPBS before further phonotypical or functional FACS analysis or cryo-preservation.

### Analysis of Cytokine Concentrations in Plasma

Blood plasma was stored at -80 °C prior to analysis. IFNγ levels were determined by LEGENDplex− from BioLegend according to the manufacturer’s instructions.

### Cell Culture

The AML cell lines HL-60 and MM6 were maintained in R10 or R20 (RPMI 1640 medium, HEPES (Gibco−), supplemented with 100 IU/ml Penicillin, 100 µg/ml Streptomycin (Gibco−) and 10 % (HL-60) or 20 % (MM6) FCS (Biochrome)).

### Phenotypical and Functional Analysis of NK Cells and T Cells

PBMCs were thawed 16 h prior to co-culture with MM6 or HL-60 cells. MM6 or HL-60 cells were either stimulated with 100 U/ml IFNγ or maintained in medium alone for 16 h. IFNγ-treated and untreated AML cells were labeled with Tag-it Violet™ Proliferation and Cell Tracking Dye (BioLegend®) and subsequently co-cultured with PBMCs for 8 h at 37 °C at an effector-to-target ratio of 10:1. As a control, AML cells and PBMCs were incubated separately in medium for the same duration. Functional assays using PBMCs from buffy coats were carried out in the presence of 10 ng/ml Brefeldin A (Sigma-Aldrich). Viability dyes and surface markers (see Supplementary Tables 2 and 3) were stained sequentially in DPBS (viability dye) or Brilliant Stain Buffer (BD Horizon™, surface markers). Intracellular markers were stained in Permeabilization Buffer following fixation with IC Fixation Buffer (both from eBioscience™). All staining steps were performed for 15 min at room temperature. Annexin V (Invitrogen) staining of AML cells was performed after viability dye staining according to the manufacturer’s instructions. For Annexin V staining, no subsequent fixation or permeabilization steps were applied.

### Statistical Analysis

Statistical analyses were performed using GraphPad Prism 10.3.0 software (GraphPad Software, San Diego California USA). For the comparison of two groups, parametric paired or unpaired t-tests were performed. More than two groups were compared by one-way ANOVA. The applied statistical tests are indicated in the figure legends.

### Study Approval

Written informed consent was obtained from all participants and the study was approved by the ethics committee of the Medical Faculty of the Heinrich-Heine-University, Germany (#2020-1222).

## Results

### CMV Reactivation in HSCT Recipients Is Associated with a Transient IFNγ Increase and Elevated Granzyme B Levels in NKG2C^pos^ NK Cells

The expansion of NK cells expressing the activating receptor NKG2C in the setting of CMV infection is well documented (reviewed in [19]). Given our hypothesis that NKG2C-positive (NKG2C^pos^) NK cells may contribute to the elimination of residual leukemic stem cells, we first quantified the frequency of NKG2C^pos^ NK cells in the bone marrow (BM) of 23 patients post HSCT, as this compartment represents the primary niche for leukemic stem cells. In line with results from peripheral blood [27], the frequency of NKG2C^pos^ NK cells was significantly increased within the BM of CMV-seropositive HSCT recipients compared to CMV-negative HSCT recipients (Figure 1 A, p = 0.0014).

**Figure 1:**
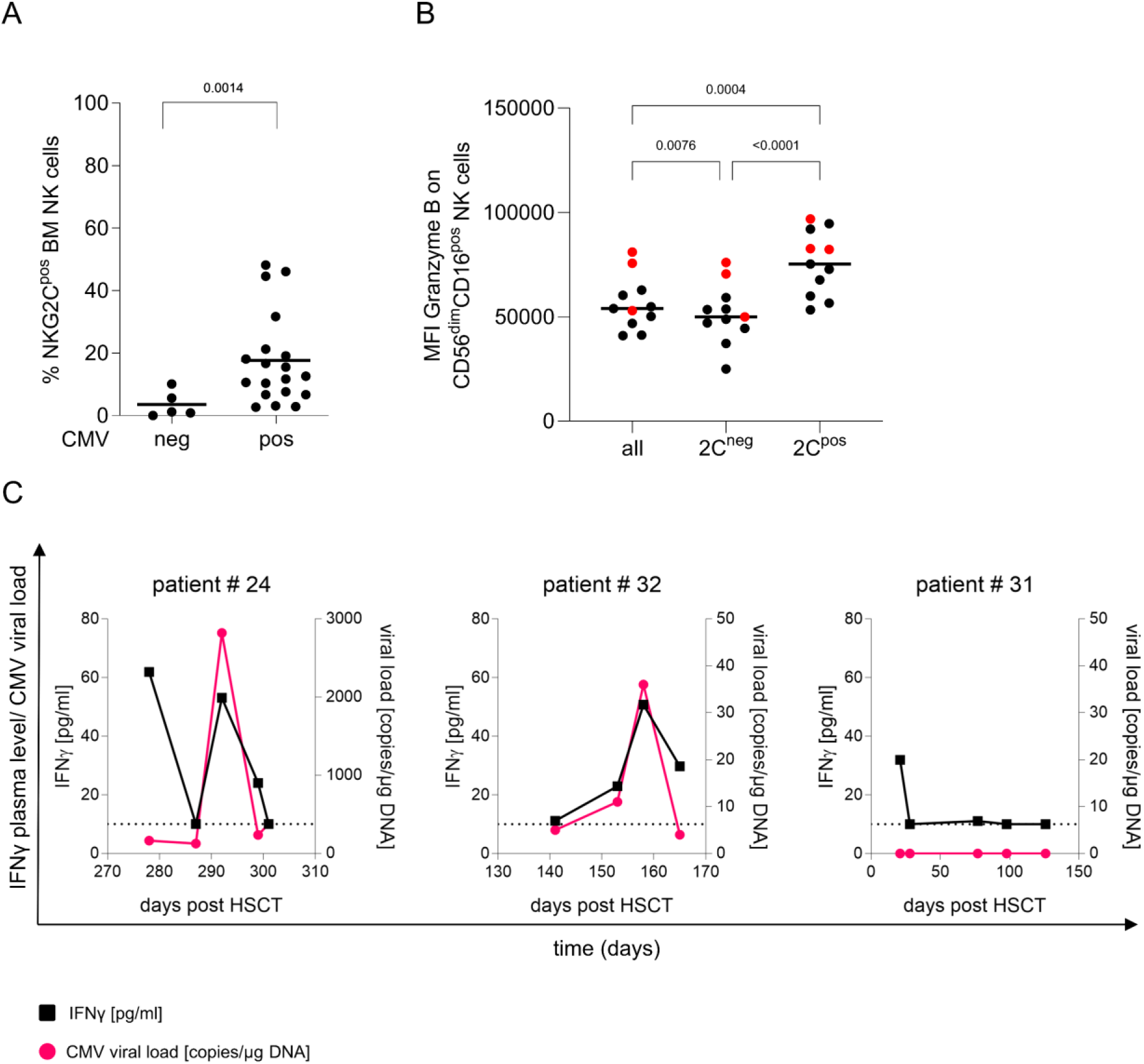
CMV Replication Is Linked to Elevated Granzyme B Levels in NKG2C^pos^ Bone Marrow NK Cells and a Temporary Increase in IFNγ Plasma Levels in Individuals Experiencing Reactivation. **A** Bone marrow MNCs were isolated from 23 AML patients that received an HSCT and NKG2C^pos^ NK cell frequencies were analyzed in the context of their CMV serostatus (Welch’s unpaired t-test). **B** Baseline Granzyme B levels were analyzed in patients with primary hematologic diseases (7 AML, 1 Non-Hodgkin Lymphoma, 1 Acute Lymphoblastic Leukemia, 2 MDS) on bulk CD56^dim^CD16^pos^ NK cells (all), as well as on NKG2C^neg^ (2C^neg^) and NKG2C^pos^ (2C^pos^) CD56^dim^CD16^pos^ NK cells (one-way ANOVA followed by Tukey’s multiple comparison). Samples from patients exhibiting CMV replication at the time of sampling are depicted as red dots. **C** Blood plasma from three AML patients was collected at different time points following HSCT and was screened for CMV viral load (red points) and IFNγ levels (black squares).

To determine the cytotoxic potential of these NKG2C^pos^ NK cells, we next examined their Granzyme B expression. As CMV viral load data and corresponding sample material from the previously analyzed BM samples were unavailable, we analyzed peripheral blood (PB) NK cells from 11 patients with underlying hematologic disease that also received HSCT. NKG2C^pos^ NK cells displayed markedly higher Granzyme B levels than their NKG2C^neg^ counterparts (Figure 1 B, p < 0.0001), an effect most pronounced in patients with detectable CMV DNAemia (red dots).

Another well-described phenomenon associated with CMV replication is a transient increase in IFNγ levels [28, 29], which has been proposed to enhance immune responses to unrelated pathogens [30, 31]. To determine whether IFNγ is released during CMV reactivation in HSCT recipients, even during the early post-transplant phase, we analyzed plasma IFNγ concentrations alongside CMV viral loads in three AML patients post HSCT within the past year (Figure 1 C). CMV DNA levels were monitored as part of routine diagnostics for CMV disease prevention. Two of these patients (patients # 24 and # 32) experienced episodes of detectable CMV DNAemia. Strikingly, in both cases, plasma IFNγ levels closely mirrored CMV DNA kinetics, increasing and declining in parallel (Figure 1 C). A third patient (patient # 31) showed no detectable CMV DNAemia and showed IFNγ levels that remained constantly low over a period of around 100 days. Similar patterns were observed in patients with other hematologic malignancies, two of whom also received HSCT (Figure S1 A). However, when considering all patients collectively, including multiple time points from the same patient, no correlation between absolute CMV viral load and IFNγ concentration was observed (Figure S1 B). This suggests that the presence of replicating CMV, rather than the magnitude of viral DNA copies in peripheral blood, triggers IFNγ release. Consistent with previous reports [28, 29], our findings indicate that CMV viremia induces a transient increase in plasma IFNγ levels during the early post-HSCT period, also within AML patients.

### Primary AML Blasts in Bone Marrow Express High Levels of the NKG2C Ligand HLA-E, Which Are Further Upregulated by IFNγ

IFNγ is known to induce the upregulation of HLA molecules on the cell surface [32], thereby enhancing target-cell susceptibility to HLA-restricted T cell-mediated lysis. This effect extends not only to classical HLA molecules but also to non-classical molecules such as HLA-E, the ligand for the activating NK cell receptor NKG2C, which we found to be expanded in the BM of CMV-seropositive AML HSCT recipients. Consistent with this, IFNγ upregulates HLA-E expression on various cell types. We therefore assessed HLA-E surface expression on primary blasts (gating of blasts shown in Figure 2 A), derived from BM (n = 6) and PB (n = 5) of either AML or MDS patients and examined how IFNγ influences this expression. Notably, blasts in the BM, the primary niche for leukemic stem cells (even in patients with formal remission status), displayed significantly higher HLA-E levels than blasts from PB (Figure 2 B, p = 0.0361). In line with previous findings [33], IFNγ further increased HLA-E expression on BM-derived myeloid blasts, (Figure 2 C, p = 0.0405), suggesting that elevated IFNγ levels during CMV reactivation may promote HLA-E upregulation on leukemic blasts.

**Figure 2:**
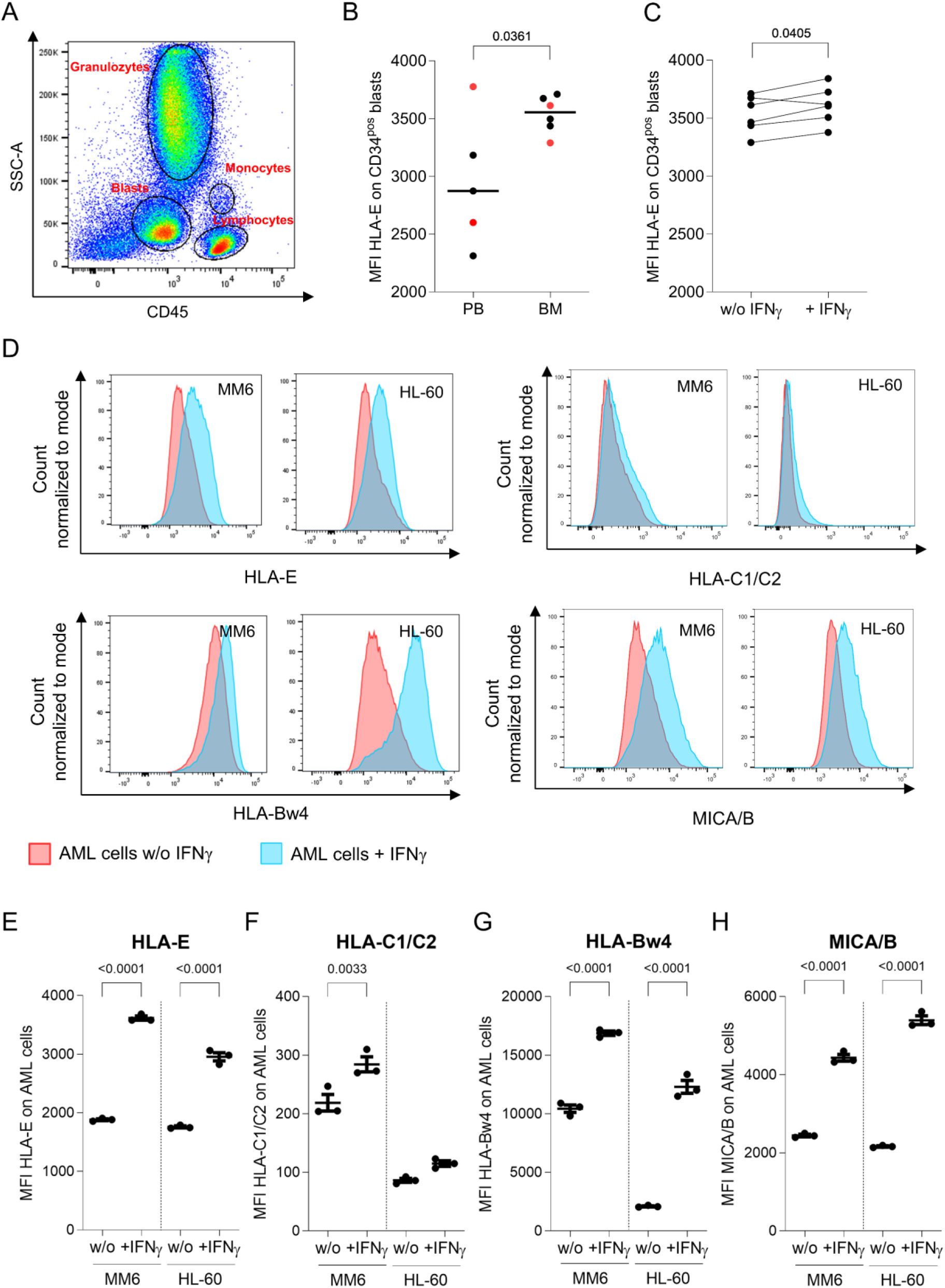
Both AML Bone Marrow Cells and Cell Lines Show High Levels of the NKG2C Ligand HLA-E After IFNγ Stimulation. **A-C** Bone marrow and peripheral blood MNCs were isolated from MDS-EB and AML patients and were allowed to rest in R10 medium or were stimulated in R10 medium supplemented with 100 U/ml IFNγ for 16 hours. HLA-E levels were analyzed on myeloid blast from samples with a minimum of 2 % blasts. **A** The gating strategy for the identification of myeloid blasts according to their relative position in a SSC-A *vs*. CD45 expression plot is shown. **B** HLA-E expression on CD34^pos^ PB and CD34^pos^ BM blasts was compared (unpaired t-test). Samples from patients with active disease are depicted in red. **C** HLA-E expression of either untreated or IFNγ stimulated CD34^pos^ BM blasts is shown (paired t-test). **D-H** MM6 and HL-60 cells were treated with 100 U/ml IFNγ for 16 hours and induction of the NK cell ligands HLA-E, HLA-Bw4 and HLA-C1/C2 were analyzed via flow cytometry. **D** Representative histograms of ligand expression of unstimulated (red histogram) and IFNγ stimulated (blue histograms) MM6 and HL-60 cells. **E-H** statistical evaluation of HLA-E (**E**), HLA-C1/C2 (**F**), HLA-Bw4 (**G**), MICA/B (**H**) expression.

To test our hypothesis that NK cells, particularly CMV-associated NKG2C^pos^ NK cells, participate in AML cell control during CMV replication, we next examined how IFNγ modulates NK cell functionality against AML targets. Due to limited availability of primary AML samples, we used the AML cell lines MM6 and HL-60 as surrogate targets and PBMCs from healthy CMV-seropositive donors as NK cell sources. IFNγ stimulation increased the expression of HLA-E as well as HLA-Bw4, HLA-C1/C2 (MM6 only) and MICA/B on both AML cell lines (Figure 2 D-H). Importantly, the magnitude of upregulation differed between the two AML cell lines, indicating that the impact on NK cell responses may vary depending on the target cell’s receptor ligand profile determined by its genetic background.

### IFNγ Increases Susceptibility of AML Cell Lines to NK Cell-Mediated Cytotoxicity

We next examined how IFNγ influences the susceptibility of AML cells to apoptosis. Annexin V staining was performed on HL-60 and MM6 cells (gating strategy in Figure 3 A) co-cultured with PBMCs from various CMV-seropositive donors, with or without prior IFNγ treatment. In the presence of PBMCs, IFNγ-pretreated MM6 and HL-60 cells showed a markedly higher frequency of Annexin V-stain positive (Annexin V^pos^) cells compared with untreated controls (Figure 3 B and 3 C, both p < 0.0001), indicating that IFNγ sensitizes AML cells to NK cell-mediated apoptosis. Among the two cell lines, MM6 cells were overall more susceptible to apoptosis than HL-60 cells (Figure S2 A), likely reflecting differences in their ability to upregulate NK cell receptor ligands, including inhibitory molecules such as HLA-Bw4. Notably, IFNγ alone induced only a mild but statistically significant increase in Annexin V^pos^ cells (0.57 % *vs*. 1.81 % in HL-60 and 1.17 % *vs*. 4.48 % in MM6; Figure S2 B), confirming that the pronounced apoptotic effects required the presence of PBMCs.

**Figure 3:**
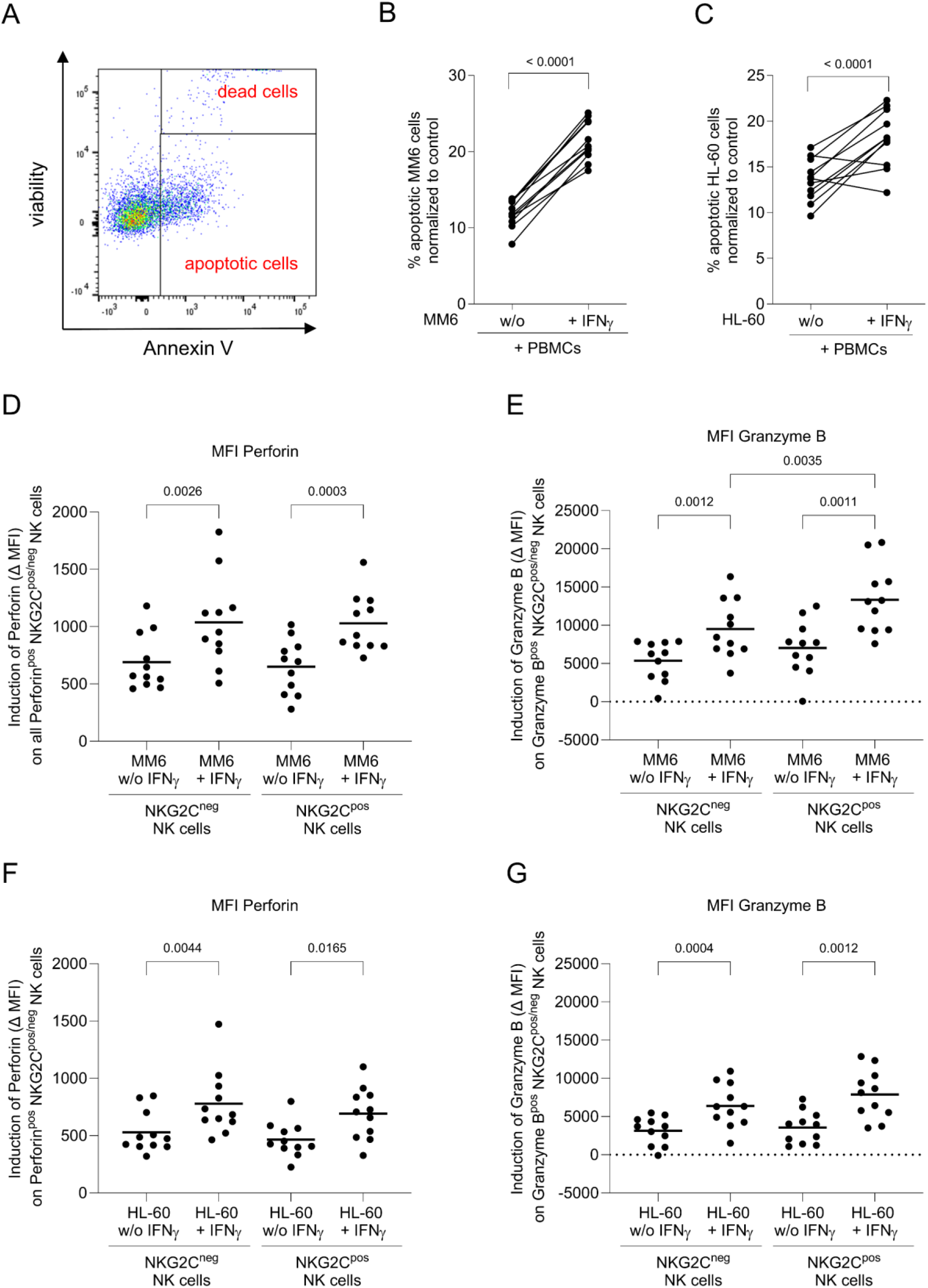
IFNγ Sensitizes AML Cells to Apoptosis and Promotes NK cells to Produce Increased Cytotoxic Molecules When Stimulated by IFNγ-Treated AML cells. The AML cell lines MM6 and HL-60 were either left untreated or stimulated for 16 h with 100 U/ml IFNγ and subsequently cocultured with PBMCs from 11 CMV seropositive healthy blood donors for 8 h. Apoptotic AML cells were determined by Annexin V staining analysis. **A** A representative Annexin V staining of HL-60 cells is shown. **B, C** Proportions of apoptotic MM6 (**B**) or HL-60 (**C**) cells induced through stimulation with PBMCs are determined by subtraction of positive cell frequencies without stimulation with PBMCs according to the respective stimulation condition. Differences are compared by paired t-tests. **D-G** Here, BFA was added to the cell culture to determine the Induction of Perforin (**D, F**) and Granzyme B (**E, G**) within NKG2C^neg^ and NKG2C^pos^ CD56^dim^ NK cells after 8 h of co-cultivation with MM6 (**D, E**) or HL-60 cells (**F, G**) that had or had not been pre-treated with IFNγ. Induction of Perforin and Granzyme B was defined as the difference of Perforin or Granzyme B MFI after stimulation with AML cells compared to baseline MFI without AML cell stimulation. Differences between groups were analyzed via one-way ANOVA followed by Tukey’s multiple comparison.

We next compared the functional responses of NKG2C^pos^ and NKG2C^neg^ NK cells by assessing Perforin and Granzyme B induction after stimulation with AML cells that had or had not been pretreated with IFNγ (Figure 3 D-G). Consistent with our earlier observation that NKG2C^pos^ NK cells in CMV-seropositive AML patients express the highest Granzyme B levels (Figure 1A), NKG2C^pos^ NK cells from CMV-seropositive PBMCs also exhibited higher baseline Granzyme B expression than NKG2C^neg^ NK cells (Figure S2 C). To quantify the Perforin and Granzyme B induction, we calculated the difference between post-stimulation mean fluorescence intensity (MFI) and baseline MFI, yielding the AML-induced expression level. Stimulation with either MM6 (Figure 3 D-E) or HL-60 cells (Figure 3 F-G) induced Perforin and Granzyme B expression in NK cells. Notably, paralleling the increased apoptosis observed in Annexin V assays, IFNγ pretreatment of MM6 and HL-60 cells significantly enhanced Perforin and Granzyme B induction in both NKG2C^pos^ and NKG2C^neg^ NK cells. Remarkably, stimulation with IFNγ-pretreated MM6 cells resulted in significantly higher Granzyme B induction in NKG2C^pos^ NK cells compared with NKG2C^neg^ NK cells (Figure 3 E, p = 0.0035), potentially reflecting the strong IFNγ-driven upregulation of HLA-E on MM6 cells. Interestingly, also NKG2C-expressing T cells were more often Granzyme B positive (S2 D) and, similar to NK cells, also showed higher Granzyme B levels compared to NKG2C^neg^ T cells (S2 E). These NKG2C^pos^ T cells upregulated Granzyme B when stimulated with IFNγ-pretreated AML targets, highlighting a broader IFNγ-mediated enhancement of immune responses across cytotoxic lymphocyte populations (Figure S2 F-G). To determine whether such NKG2C-expressing T cells are present in the bone marrow of HSCT recipients, we reanalyzed our BM data and quantified NKG2A and NKG2C expression within the (CD3/CD19)^high^ compartment, which predominantly contains T cells (see S2 H for gating). Similar to the elevated NKG2C^pos^ NK cell frequencies, we observed increased frequencies of NKG2C^pos^ (CD3/CD19)^high^ cells in the bone marrow (Figure S2 I-J), suggesting that these T cells may also contribute to functional interactions within the NKG2C-HLA-E axis.

## Discussion

Although accumulating evidence indicates that CMV replication reduces relapse risk in AML patients undergoing HSCT, the underlying mechanisms remain incompletely understood. Koldehoff *et al*. proposed a direct pro-apoptotic effect of CMV on leukemia cells [34]. However, while *in vitro* infection rates of myeloid progenitor cells can exceed 90 %, the frequency of natural CMV infection *in vivo* is substantially lower, with only ∼0.01-0.001 % of progenitor cells expressing detectable latency-associated transcripts [35]. This pattern is characteristic of a largely transcriptionally silent, latent infection rather than productive viral replication. Reactivation mainly occurs during terminal myeloid differentiation [35], a developmental stage typically not reached by AML blasts due to their differentiation arrest (reviewed in [36]).

Interestingly, beneficial effects of CMV on relapse risk have also been observed in individuals without detectable CMV DNAemia [37], suggesting that durable CMV-driven immunological imprints may contribute to protection. Several groups have hypothesized a role for NKG2C^pos^ NK cells in this process (reviewed in [24]). We propose that the protective effect arises not from NKG2C^pos^ NK cells alone but from the combination of NKG2C^pos^ NK cells and elevated IFNγ levels during CMV reactivation. Transient IFNγ increases following CMV infection have been shown to enhance immunity, including in the context of influenza vaccination [30]. Here, we demonstrate that HSCT recipients with CMV viremia exhibit a temporary rise in plasma IFNγ concentrations that closely parallels viral DNA levels and returns to baseline once viremia resolves. This observation may explain why active CMV replication, rather than CMV seropositivity alone, is required to confer protection. In our experiments, IFNγ treatment alone increased AML cell apoptosis both in the presence and absence of NK cell-containing PBMCs from CMV-seropositive individuals. Moreover, the CMV major immediate-early promoter-enhancer contains two functional gamma interferon-activated site-like elements and IFNγ has been shown to activate immediate early gene expression in latently infected progenitor cells [35, 38], potentially triggering viral reactivation and facilitating the elimination of CMV-infected AML cells through lytic replication.

In addition to the direct effects of transient IFNγ accumulation on AML cells, we examined how NK cells, which are among the earliest lymphocyte populations to reconstitute after HSCT, contribute to enhanced leukemic stem cell killing. Since NK cell activation is dependent on the balance of activating and inhibitory signals, we analyzed the expression of HLA molecules on IFNγ-stimulated AML cells, as these serve as key ligands for NK cell receptors. Notably, IFNγ induced robust upregulation of HLA-E on primary AML blasts as well as on AML cell lines (HL-60, MM6). Given that HLA-E is the ligand for the activating NK cell receptor NKG2C, CMV reactivation may indirectly enhance NKG2C^pos^ NK cell responses, consistent with our observation that these cells are enriched in the bone marrow of CMV-seropositive AML patients. Moreover, both the CMV-encoded UL40 signal peptide and signal peptides derived from IFNγ-induced elevated HLA class I expression can further stabilize HLA-E on the cell surface. This suggests that IFNγ may promote HLA-E upregulation not only in CMV-infected AML cells but also in uninfected ones [39-41]. In this context, the HLA-E genotype (HLA-E*01 *vs*. *03), along with CMV strain-specific UL40 peptide variants, may modulate the strength of NKG2C^pos^ NK cell activation, either enhancing or dampening effector responses [41-45]. In addition, IFNγ upregulated the activating NKG2D ligand MICA/B, providing a plausible explanation for why NKG2C^neg^ NK cells also exhibited enhanced Perforin and Granzyme B induction in the presence of IFNγ.

While HLA-Bw4 and HLA-C1/C2, both of which were also upregulated by IFNγ, exclusively deliver inhibitory signals to NK cells through engagement of inhibitory KIRs, HLA-E is unique in that it can transmit either inhibitory or activating signals depending on whether NK cells express NKG2A or NKG2C. Because NKG2A^pos^ NK cells constitute the dominant subset in CMV-seronegative individuals, and only a small fraction of NK cells co-express both receptors, HLA-E is generally perceived as an inhibitory ligand, restraining NK cell activity via NKG2A.

Indeed, blockade of the NKG2A-HLA-E axis is under active clinical investigation as an immunotherapeutic strategy, including in AML and MDS patients undergoing haploidentical transplantation (NCT06892223). Monalizumab, a humanized anti-NKG2A antibody, disrupts NKG2A-HLA-E interactions and thereby enhances both NK- and T cell-mediated antitumor immunity. Although a phase III trial of Monalizumab plus Cetuximab in previously treated head and neck squamous cell carcinoma did not improve overall survival relative to Cetuximab alone [46], promising response rates in earlier phase II studies underscore the therapeutic potential of NKG2A blockade while highlighting the need for deeper mechanistic insight [47]. Unlike many classical HLA class I molecules, which are often downregulated by tumor cells to evade cytotoxic T lymphocytes, HLA-E frequently remains stably expressed and can even be upregulated, in a variety of solid tumors and lymphomas [48-50]. Consistent with this, we observed higher HLA-E expression on bone marrow-resident myeloid blasts compared with blasts in peripheral blood. Furthermore, NKG2A is expressed at high frequencies on NK cells from AML patients [51] and previous studies have shown that IFNγ-driven HLA-E upregulation on AML blasts can impair NKG2A-dependent NK cell cytolysis following haploidentical HSCT [33].

In contrast to the inhibitory effects mediated through NKG2A, expansion of NKG2C^pos^ NK cells has been associated with lower rates of AML relapse after HSCT [52]. In our study, we show that NK cells produce substantial amounts of the cytotoxic effector molecules Perforin and Granzyme B and that these responses are further enhanced when AML target cells are pretreated with IFNγ. This indicates that NK cells are highly “armed” against AML cells under inflammatory conditions. Notably, when stimulated with IFNγ-pretreated MM6 cells, NKG2C^pos^ NK cells produced significantly higher levels of Granzyme B than their NKG2C^neg^ counterparts, underscoring a synergistic interaction between NKG2C^pos^ NK cells and IFNγ in this context. Importantly, when analyzing HSCT recipients, we found that NKG2C^pos^ NK cells expressed the highest Granzyme B levels, suggesting that these cells may be particularly well equipped to eliminate residual leukemic stem cells during the early post-transplant period when tumor-specific T cells are largely absent. Unfortunately, the limited availability of patient material prevented us from further dissecting the cytotoxic potential of primary NKG2C^pos^ NK cells. We also attempted to interrogate the NKG2C-HLA-E axis more directly by blocking NKG2C before AML stimulation; however, NKG2C antibodies themselves activated NK cells, precluding unbiased analysis. Nevertheless, despite the transient nature of both NKG2C^pos^ NK cell activation and IFNγ elevation during CMV viremia, HSCT recipients who experience CMV replication likely benefit from durable immune remodeling. Highly functional NKG2C^pos^ NK cells have been shown to undergo long-term expansion in HSCT recipients following CMV viremia [20, 53, 54], suggesting that these cells may continue to contribute to AML control even at later time points.

Interestingly, the protective effect of CMV viremia after HSCT in AML patients is lost when either T cells or NK cells are depleted *in vitro* or *in vivo* [5, 8, 55], indicating that T cells also contribute to this phenomenon. In our study, we observed that NKG2C-expressing T cells upregulated Granzyme B production when stimulated with IFNγ-pretreated AML cells, despite not being specific for MM6 or HL-60 antigens. This highlights a potentially important, antigen-independent role for T cells in AML control. A more detailed analysis of T cell involvement would be of considerable interest. Foley *et al*. previously reported that NK cell function is impaired in HSCT recipients when T cells are depleted, suggesting that T cells may support NK cells in mediating protection against AML relapse [14]. Additional evidence from umbilical cord blood transplant studies has shown that NKG2C^pos^ CD8 T cells expand in response to CMV reactivation. Similarly, NKG2C^pos^ CD56^high^ CD161^neg^ CD8 T cells have been identified in the liver, closely resembling the NKG2C^pos^ CD8 T cells described in cord blood recipients. These cells exhibit a restricted TCR repertoire and display transcriptional and functional profiles that are highly reminiscent of NK cells, particularly following NKG2C ligation or cytokine stimulation. Importantly, these T cells are not specific for CMV pp65 but are predominantly HLA-E-restricted, enabling their activation through both NKG2C and HLA-E whose expression is further enhanced during CMV reactivation in response to IFNγ [56, 57]. HLA-E-restricted CD8 T cells may therefore play a crucial role in pathogen and tumor control in both HLA-matched and mismatched donor-recipient settings, exploiting the NKG2C-HLA-E axis. This may be particularly relevant when classical HLA class I molecules are downregulated, as frequently observed in cancer and chronic infection (reviewed in [45, 50, 58]).

For potential clinical translation, NK cell-based therapies represent an attractive strategy, as NK cells have demonstrated strong antitumor activity with minimal adverse effects (reviewed in [59]). Liu *et al*. showed that *in vitro*-expanded NKG2C^pos^ NK cells are highly effective killers of allogeneic pediatric T and precursor B cell ALL blasts [60]. Interestingly, this cytotoxicity was NKG2C-independent due to the absence of HLA-E on the target cells and instead appeared to be mediated by KIR interactions. However, considering the IFNγ-induced upregulation of HLA-E on primary AML blasts observed in our study, we propose that adoptive transfer of NKG2C^pos^ NK cells, particularly when combined with IFNγ treatment, may represent a promising therapeutic approach for AML. Such strategies could be further potentiated by immune checkpoint blockade, for example using PD-1/PD-L1 inhibitors or Monalizumab. Taken together, our findings support a model in which NKG2C^pos^ NK cells, NKG2C^pos^ CD8 T cells and IFNγ synergize to mediate the anti-leukemic effects clinically associated with active CMV replication.

## Acknowledgements

We thank Nicole Bombis, Stefanie Geyh, Irmgard Hamann, Saskia Mayer and all participating physicians for their assistance in sample collection and preparation and all patients for participating in the study. This work was supported by the Düsseldorf School of Oncology (funded by the Comprehensive Cancer Center Düsseldorf/Deutsche Krebshilfe and the Medical Faculty HHU Düsseldorf).

---

**Figure S1:**
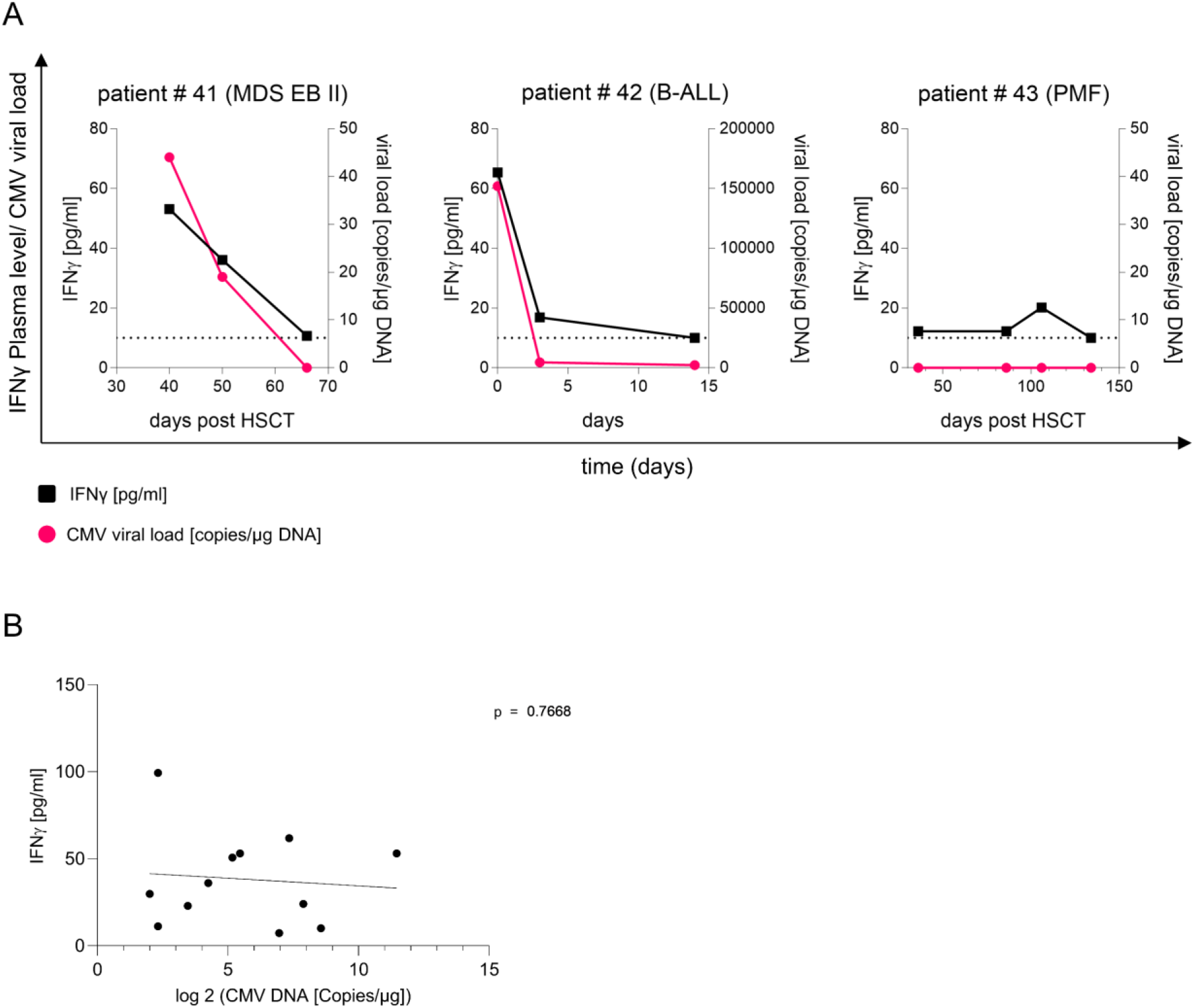
**A** Blood plasma from three patients with underlying hematologic disease (Myelodysplastic Syndromes with Excess of Blast II (MDS EB II), B cell ALL (B-ALL), and Primary myelofibrosis (PMF) with (patient # 41 and # 43) or without HSCT (patient # 42) was collected at different time points and was screened for CMV viral load (red points) and IFNγ levels (black squares). **B** Correlation of IFNγ plasma levels and CMV viral load of AML (n = 2) or MDS (n = 2) patients with active CMV replication at the time of sampling is depicted.

**Figure S2:**
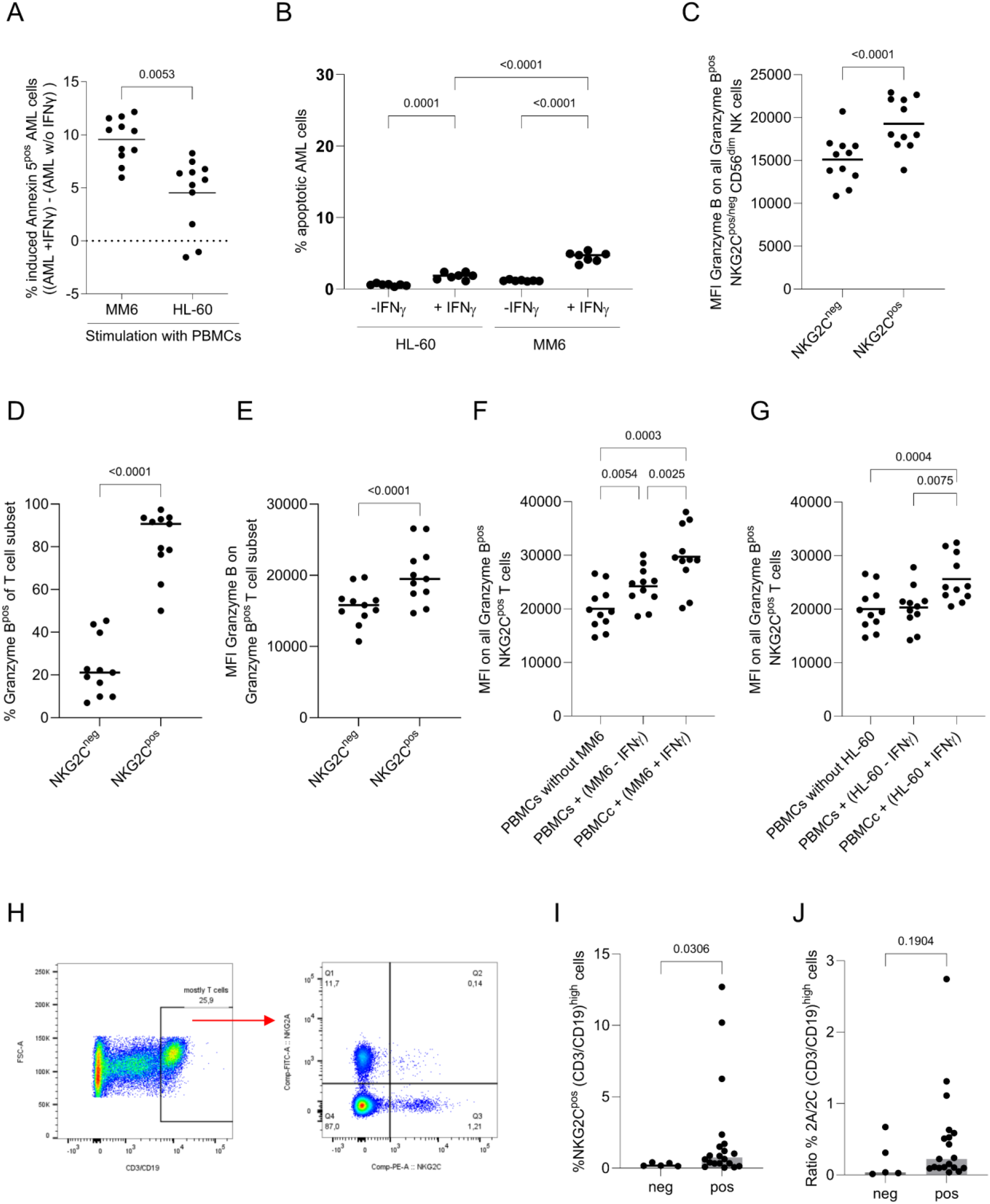
**A, B** The AML cell lines MM6 and HL-60 were left either untreated or stimulated for 16 h with 100 U/ml IFNγ and subsequently cocultured with PBMCs from 11 CMV seropositive donors for 8 h. **A** Comparison of induced apoptotic MM6 vs HL-60 cells upon stimulation with PBMCs from CMV seropositive healthy individuals is shown. Apoptotic AML cells were determined by Annexin V stain analysis and induction was calculated as the difference of apoptotic IFNγ-stimulated and -unstimulated AML cells after co-incubation with PBMCs (paired t-test). **B** Proportions of apoptotic MM6 or HL-60 cells induced at baseline levels (-IFNγ) or through stimulation with IFNγ (+IFNγ) are determined via Annexin V staining (n = 7). Differences are compared by one-way ANOVA followed by Šidák’s multiple comparisons. **C** Granzyme B baseline levels of NKG2C^pos^ and NKG2C^neg^ NK cells from CMV seropositive healthy blood donors are shown (paired t-test). **D**,**E** Granzyme B^pos^ T cell frequencies (**D**) and Granzyme B Levels (**E**) within NKG2C^neg^ and NKG2C^pos^ T cells from healthy donors are shown. **F**,**G** Granzyme B expression within NKG2C^pos^ T cells in dependency of stimulation with MM6 (**F**) or HL-60 (**G**) with the addition of BFA is shown (one-way ANOVA, followed by Tukey’s multiple comparison). **H** Gating strategy for identification of NKG2A^pos^ and NKG2C^pos^ (CD3/CD19)^high^ “T cells” within bone marrow samples. **I** Comparison of NKG2C^pos^ (CD3/CD19)^high^ “T cell” frequencies between CMV seronegative and seropositive HSCT recipients. **J** To control for potential contributions from B cells, CD4 T cells and CD4/CD8 double-negative T cells, the ratio of NKG2A^pos^ to NKG2C^pos^ cells was determined. Differences between CMV negative and positive individuals in I and J were analyzed via Welch’s unpaired test.

